# Nucleosome condensate and linker DNA alter chromatin folding pathways and rates

**DOI:** 10.1101/2024.11.15.623891

**Authors:** Yunrui Qiu, Shuming Liu, Xingcheng Lin, Ilona Christy Unarta, Xuhui Huang, Bin Zhang

## Abstract

Chromatin organization is essential for DNA packaging and gene regulation in eukaryotic genomes. While significant progresses have been made, the exact atomistic arrangement of nucleosomes remains controversial. Using a well-calibrated residue-level coarse-grained model and advanced dynamics modeling techniques, particularly the non-Markovian dynamics model, we map the free energy landscape of tetra-nucleosome systems, identify both metastable conformations and intermediate states in folding pathways, and quantify the folding kinetics. Our findings show that chromatin with 10*n* base pairs (bp) DNA linker lengths favor zigzag fibril structures. However, longer linker lengths destabilize this conformation. When the linker length is 10*n* + 5 bp, chromatin loses unique conformations, favoring a dynamic ensemble of structures resembling folding intermediates. Embedding the tetra-nucleosome in a nucleosome condensate similarly shifts stability towards folding intermediates as a result of the competition of inter-nucleosomal contacts. These results suggest that chromatin organization observed *in vivo* arises from the unfolding of fibril structures due to nucleosome crowding and linker length variation. This perspective aids in unifying experimental studies to develop atomistic models for chromatin.

**Significance:** Atomic structures of chromatin have become increasingly accessible, largely through cryo-EM techniques. Nonetheless, these approaches often face limitations in addressing how intrinsic *in vivo* factors influence chromatin organization. We present a structural characterization of chromatin under the combined effects of nucleosome condensate crowding and linker DNA length variation—two critical *in vivo* features that have remained challenging to capture experimentally. This work leverages a novel application of non-Markovian dynamical modeling, providing accurate mapping of chromatin folding kinetics and pathways. Our findings support a hypothesis that *in vivo* chromatin organization arises from folding intermediates advancing toward a stable fibril configuration, potentially resolving longstanding questions surrounding chromatin atomic structure.

## Introduction

Chromatin organization, essential for packaging DNA in eukaryotic genomes, plays a crucial role in various genetic functions.^1–5^ Atomistic structures of chromatin, where available, have proven invaluable in constructing mechanistic models of gene regulation and other processes.^6,7^ However, the detailed organization at the atomic level has been the subject of contentious debate. The existence of 30-nm fibers, characterized by nucleosomes following a zig-zag path and stacking on top of each other to form a twisted two-column fibril, has long been documented.^6–10^ However, several *in vivo* experimental techniques, including cryo-electron microscopy,^6^ Micro-C,^11^ and ChromEMT,^12^ have indicated a lack of ordered fibril-like structures. Instead, these techniques suggest the presence of 10-nm disordered arrays with prevailing local oligomer motifs, such as trimers, *α*-tetrahedron, and *β*-rhombus tetramers.^11–16^ These contradictory observations have left the folding principles of chromatin, particularly at the short length scale of tens of nucleosomes, unresolved.

A more dynamic perspective on chromatin organization has proven to be insightful. Ding et al. ^17^ characterized the folding landscape of a tetra-nucleosome using a residue-level coarsegrained (CG) model, proposing that the unfolding of fibril structures leads to the irregular conformations observed in the nucleus, thus bridging *in vitro* and *in vivo* configurations. Liu et al. ^18^ further demonstrated that unfolding can generate clutches, consistent with observations from super-resolution imaging of the cell nucleus.^19^ A recent cryo-electron tomography (cryo-ET) study further supports the presence of similar folding principles for chromatin both *in vitro* and *in situ*.^12^ However, the computational studies employed biased simulations to accelerate chromatin folding and unfolding. While thermodynamic predictions can be made, extracting accurate kinetic information from such simulations proves challenging. Consequently, it is significant to more directly determine whether the irregular *in vivo* chromatin configurations represent intermediate folding stages leading to the zigzag fibril structure.

Markov state models (MSMs) offer an effective approach to extract interpretable kinetic information, facilitating the prediction of transition rates and the identification of reaction pathways from the molecular dynamics (MD) data.^20–35^ Specifically, MSMs could coarse-grain complex MD configurations into conformational states and model continuous dynamics as Markovian jumps between these states at discrete time interval (i.e., lag time). The construction of such models and prediction of long-timescale dynamics require only distributed, relatively short MD trajectories as inputs, which can be efficiently generated in parallel using supercomputing facilities. To aid the interpretation of complex dynamics, it is often beneficial to build a model with only a handful of metastable and interpretable states. However, accurately capturing the slow dynamics between these states is challenging, as this requires a Markovian lag time that is typically much longer than the length of the affordable MD simulation. Our recently developed non-Markovian dynamic model demonstrates significant advantages in addressing this challenge. By employing the generalized master equation (GME) with memory kernel functions, it could effectively model non-Markovian transition dynamics, greatly enhancing modeling accuracy.^36–39^

In this study, we conduct extensive residual-level CG MD simulations to examine chromatin folding. Specifically, we investigate the folding of four systems: three isolated tetranucleosomes with varying nucleosome repeat lengths (NRL) of 167, 172, and 177 base pairs (bp), and a tetra-nucleosome with an NRL of 167 bp embedded within a nucleosome condensate. We construct microstate-MSM and interpretable non-Markovian dynamics model for each system to comprehensively characterize the folding pathways and kinetics. In every system, transition path analysis consistently reveals numerous parallel folding pathways with comparable fluxes. This finding contrasts with typical protein folding and more closely resembles multi-body self-assembly processes. Notably, intermediate states along these pathways resemble *in vivo* chromatin organization. Furthermore, while the tetra-nucleosome with NRL = 167 bp folds stably into the zigzag fibril structure, both the nucleosome condensate and a 5 bp longer DNA linker destabilize the fibril conformation, promoting the formation of folding intermediates through different molecular mechanisms. Extending the linker DNA length by 10 or 5 bp has markedly different effects on the chromatin folding landscape: while the 10 bp longer linker favors the zigzag fibril configuration, the 5 bp longer linker does not result in any specific preferred structure, instead favoring an ensemble of dynamic conformations. These results reinforce the idea that the absence of regular fibril configurations inside cell nuclei could result from chromatin unfolding driven by various factors such as crowding environments and linker length variation.

## Results

### Construction of dynamics models for chromatin folding from extensive MD simulations

Our study focuses on tetra-nucleosomes, the fundamental units of a zigzag fibril configuration,^6,7^ to explore chromatin organization in various biological contexts. Specifically, we constructed a series of tetra-nucleosome structures with 20, 25, and 30 bp linkers, corresponding to NRL of 167, 172, and 177 bp (Figure 1A-C). We define nucleosomal DNA as 147 bp, and refer to the DNA connecting two adjacent nucleosomes as the linker DNA. To simulate a biologically relevant scenario, we further embeded the tetra-nucleosome with the 20 bp linker within a solution of single nucleosomes to account for nuclear crowding effects. Our setup results in an overall nucleosomal concentration of 0.3 mM (Figure 1D), consistent with concentrations estimated from *in vitro* nucleosome array condensates.^40,41^

**Figure 1:**
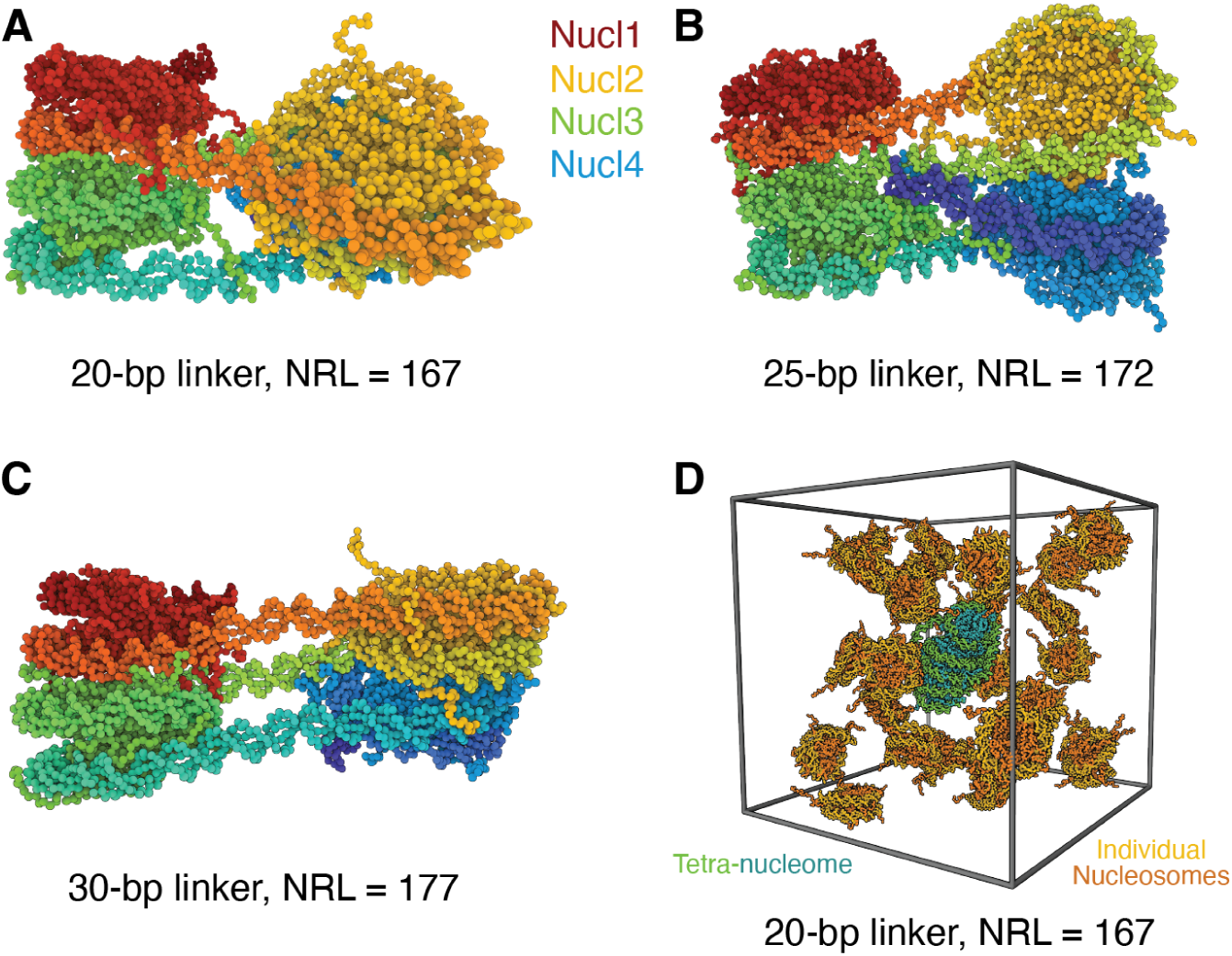
Representative configurations for the four tetra-nucleosome systems studied. The three isolated systems feature tetra-nucleosomes of 20-bp (A), 25-bp (B), and 30-bp (C) DNA linker. The corresponding NRL is 167, 172, and 177 bp, respectively. In the fourth system, the tetra-nucleosome with NRL=167 is embedded into a nucleosome condensate.

We performed MD simulations to investigate conformational dynamics of chromatin with a residue-level CG model. Coarse-graining is essential because all-atom simulations are computationally prohibitive for extensive sampling of multiple nucleosome structures. ^42–46^ Specifically, we employed a one-bead-per-amino-acid model^47^ and a three-bead-per-nucleotide model^48^ to maintain sufficient resolution for an accurate description of specific proteinprotein and protein-DNA interaction with physical chemistry potentials. These models have been effectively utilized to study various protein-DNA systems, accurately reproducing experimental results and providing mechanistic insights. ^17,18,49–54^ Further details on the CG force field are provided in the *Methods* section.

For each of these four systems, we conducted multiple independent, unbiased MD simulations to explore various regions of the chromatin configurational space. As illustrated in Figure 2A, these simulations were initiated from distinct conformations, capturing various degrees of chromatin compaction and folding, as identified in a previous study.^17^ For isolated tetra-nucleosomes with varying linker lengths, we executed 4,643 simulations for each system. In the case of the condensate system, due to its longer equilibration time, we reduced the number of trajectories to 530 and extended each simulation length. All simulations are sufficiently long to allow for the relaxation of chromatin conformations and the attainment of local equilibrium (Figures S1 and S2).

**Figure 2:**
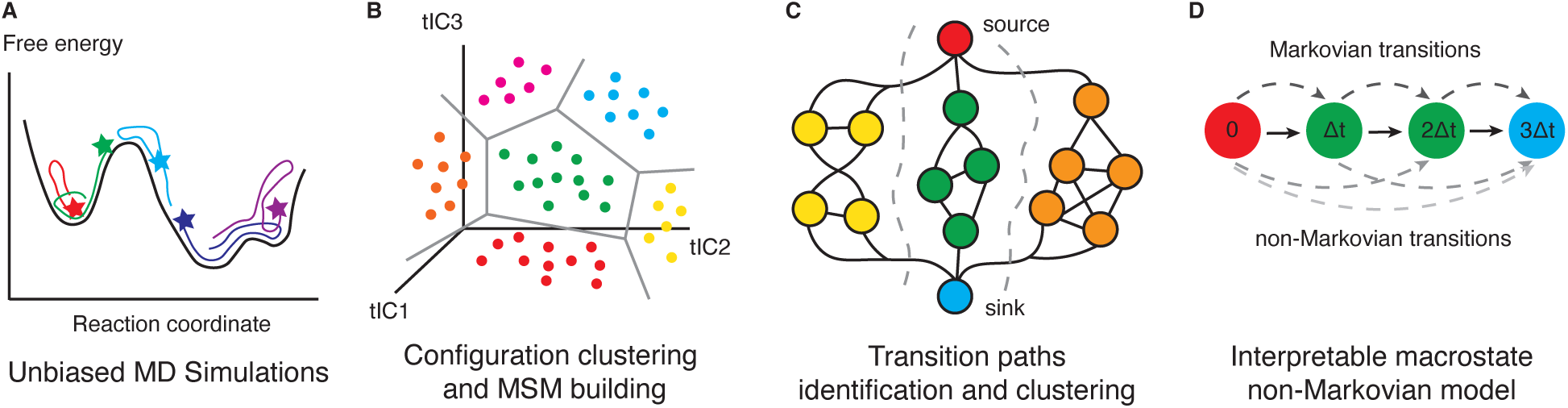
Overview of the computational pipeline for determining chromatin folding kinetics and pathways. The process begins with extensive unbiased MD simulations, initiated from a variety of configurations (A). Configurations collected from these simulations are projected onto collective variables identified by tICA and then clustered into microstates to construct MSMs. (B). Subsequently, chromatin folding pathways and reaction channels are identified using transition path theory and a path clustering algorithm (C). Finally, the microstates are lumped into a few metastable macrostates, and the transition dynamics between these macrostates are modeled using the generalized master equation that incorporates time-dependent memory kernels (D).

Subsequently, we constructed dynamics models for each system to study chromatin folding over timescales much longer than individual MD simulation trajectories. We projected tetra-nucleosome configurations explored by unbiased trajectories onto collective variables using time-lagged independent component analysis (tICA),^55–57^ and clustered them into microstates by the K-Means algorithm (Figure 2B and Figures S3-S6). Microstate-MSMs were then constructed and validated to model chromatin folding dynamics (Figures S7-S10). Further empolying the transition path theory (TPT)^20,21,58^ and path clustering algorithm,^27^ we identified the kinetic pathways and reaction channels for tetra-nucleosome folding. (Figure 2C and Figures S11-S17).

To facilitate the interpretation of folding dynamics, we also built non-Markovian dynamics models with six metastable macrostates lumped from microstates using the Integrated Generalized Master Equation (IGME) method^36^ (Figure 2D and Figures S18-S25). Unlike microstate-MSMs, which typically require hundreds of states to capture the slowest dynamics and are thereby challenging to interpret, IGME models can accurately recover the slowest dynamics with only a few metastable states. This advantage arises from IGME’s incorporation of the time-integration of memory kernels to account for non-Markovian dynamics. We demonstrated that for all systems reported in this study, IGME models consistently show significantly higher accuracy in modeling long-term dynamics compared to MSMs constructed with similar or longer MD segments. More details regarding system setup, simulations, and constructions of MSMs and IGME models are available in the *Methods* section and *Supporting Information*.

We note that CG models are frequently limited in their ability to accurately represent the timescales of dynamical motion. Nevertheless, their demonstrated accuracy in quantifying protein-DNA interactions and mapping the free energy landscape of nucleosome unwinding and chromatin folding^17,18,59–61^ suggests that they are well-suited to capture the sequence of events occurring over slow timescales, governed by the topology of the energy landscape. This capability to model the qualitative mechanisms driving large-scale conformational changes supports the comprehensive kinetic analyses presented above.

### Elucidation of tetra-nucleosome folding pathways and kinetics

We initiated our investigation with the isolated NRL = 167 tetra-nucleosome. As shown in Figure 3A, we first plotted the two-dimensional free energy landscape projected onto internucleosomal distances, *d*_13_ and *d*_24_, based on the equilibrium populations obtained from the microstate-MSM. The resulting landscape, featuring an approximate 10 kcal/mol difference between unfolded and folded regions, indicates the stability of the native tetra-nucleosome structures and is consistent with previous estimations from neural network fitting of mean forces.^17^

**Figure 3:**
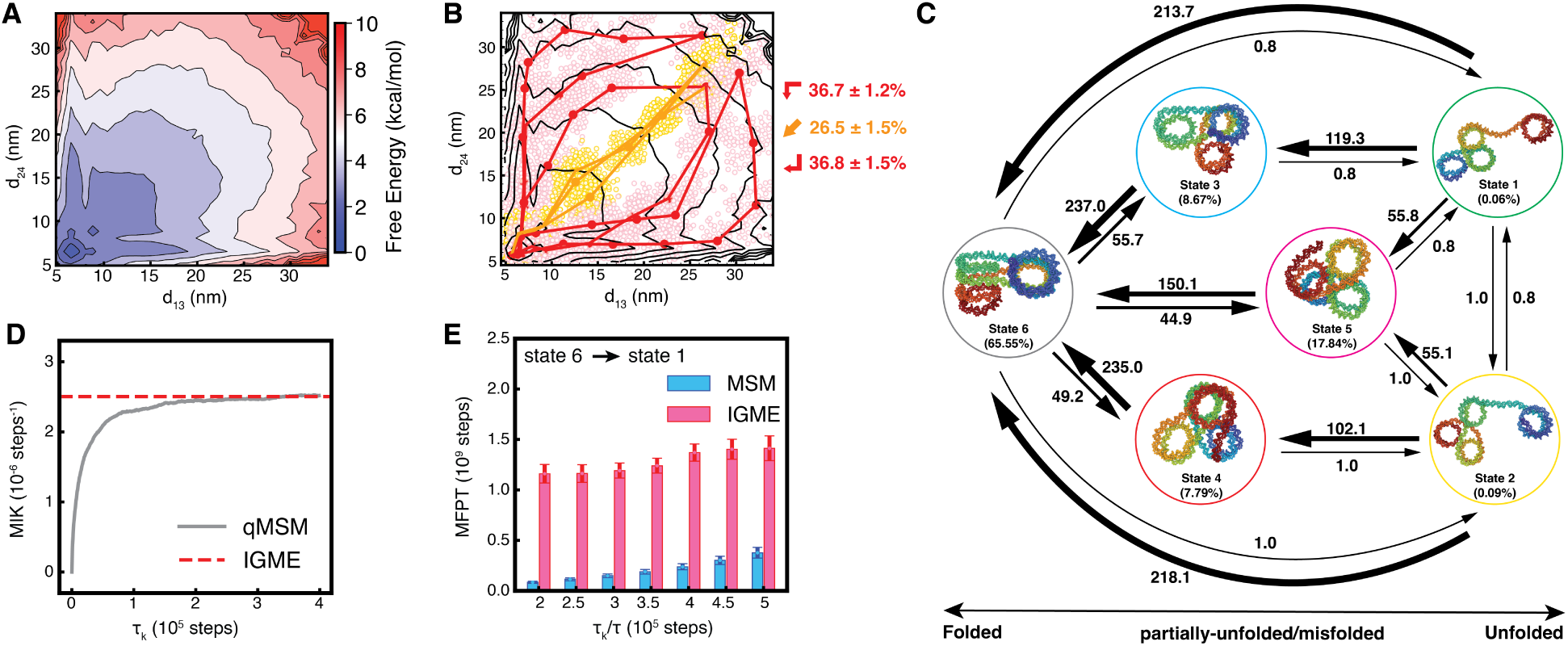
Folding pathways and kinetics for the NRL = 167 tetra-nucleosome. (A) The free energy profile along the center-of-geometry distance between 1-3 (*d*_13_) and 2-4 (*d*_24_) nucleosomes. (B) The three reaction channels for tetra-nucleosome folding. Top three transition pathways with most reactive flux from each one reaction channel are drawn as lines. The filled and open circles represent the centers of, and the MD configurations belonging to, the microstates along the pathways, respectively. The total reactive flux of each reaction channel is provided on the side. (C) Diagram of the IGME model with six macrostates. The numbers represent rates estimated as inverse MFPT labeled in unit of (10^9^ steps)*^−^*^1^ for the corresponding transitions. Histone proteins are not shown in the representative configurations of each state. (D) The mean integral of memory kernels (MIK) across different *τ_k_* for the six-macrostate model, calculated using the quasi-MSM (qMSM) and IGME. (E) The mean first passage time (MFPT) from State 6 to State 1, computed from MSMs and IGME with different lag times, is compared. The error bars show standard deviations, estimated from bootstrapping of data 20 times with replacement.

By conducting transition path analysis with the microstate-MSM, we identified over 20,000 pathways connecting the unfolded to folded states. Strikingly, the most dominant pathway only contributes 0.09% of the total reactive flux for the folding. This contrasts with typical protein folding, such as NTL9 folding, where the top 10 pathways can account for 25% of the total flux.^29^ In the case of tetra-nucleosome folding, approximately 600 pathways are required to reach the same flux level (Figure S13A). The observation of the downhill landscape and numerous parallel pathways with comparable fluxes suggests that tetra-nucleosome folding is more akin to heterogeneous aggregation in self-assembly systems than to typical protein folding.^32^

To aid in interpreting the multiple parallel pathways with similar fluxes, we grouped the pathways into a small set of metastable reaction channels using the Latent-space Path Clustering algorithm.^27^ This method identified three metastable reaction channels: two sequential channels (up and down sequential channels) and one concerted channel (Figure 3B and S17A). These reaction channels exhibit distinct folding mechanisms: the sequential channels correspond to processes where one pair of nucleosomes stacks before the other, while the concerted channel represents processes where both pairs of nucleosomes stack simultaneously. These findings align with a previous study that characterized tetra-nucleosome folding pathways using the string method. ^17^ The overall reactive flux in the sequential channels is slightly higher than in the concerted channel, indicating a preference for reactions. This preference may result from more stable intermediates encountered along sequential pathways with increased internucleosomal contacts.

To further understand chromatin folding dynamics, we constructed an IGME model with six metastable macrostates (Figure 3C). The sparsely populated states 1 and 2 correspond to unfolded structures. Their high transition rates toward the folded state (state 6), coupled with low rates for backward transitions, align with a downhill free energy landscape. States 3 and 4 present partially unfolded structures, where one nucleosome extrudes from the other three. These configurations resemble those observed in sequential chromatin folding pathways (Figure S27). A cryo-EM study directly observed tri-nucleosome configurations,^62^ demonstrating the IGME model’s capability to identify metastable states.

The model also identifies a misfolded state (state 5), marked by non-canonical stacking between the first and fourth nucleosomes (i.e., *i* & *i* + 3 contacts). For the transition to the zigzag conformation of state 6, these interactions must be disrupted, resulting in slow transition rates. We classify this state as misfolded, as transition path analysis indicates that none of the top 3,000 most reactive pathways pass through it. Future cryo-EM studies may clarify whether this state can be resolved.

Specifically, we found that our IGME model provides significant improvements over MSMs in accurately quantifying equilibrium populations and transition rates among metastable states using shorter MD simulation segments, as the memory kernel’s relaxation timescale is generally much shorter than the Markovian lag time. By evaluating memory kernel integrations over various lag times using different approaches (see *Methods* for more details), we observed that the integrations converge at approximately *τ_k_* = 2 × 10^5^ steps (Figure 3D), beyond which the IGME models can be constructed. An example IGME model built from MD simulation segments of ∼ 3 × 10^5^ steps has been shown to accurately predict dynamics compared to MD simulations (Figure S19). Conversely, the MSM constructed with a longer lag time exhibited substantially higher prediction errors, failed the Chapman–Kolmogorov test, and predicted much faster state relaxation dynamics (Figures S18–S19). Across a wide range of lag times, IGME models consistently outperformed MSMs in capturing the slowest mean first passage time (MFPT) for unfolding dynamics (Figure 3E). Matching the performance of the IGME model would require constructing the MSM with significantly longer lag times, exceeding the length of our MD simulations.

### Nucleosome condensate promotes chromatin unfolding

Our study on the isolated tetra-nucleosome does not consider the nuclear environment. The presence of additional nucleosomes may not only act as a crowding agent but also affect the stability of chromatin conformations through direct interactions. Inter-nucleosomal interactions could destabilize the chromatin fiber by forming interdigitated configurations.^18,63^ In this section, we examine the effects of immersing the tetra-nucleosome in a nucleosome condensate environment on chromatin folding.

In simulations of this condensate system, individual free nucleosomes were initially uniformly distributed throughout the cubic simulation box with the tetra-nucleosome at the center (Figure 4A). Over time, they rapidly and spontaneously aggregated around the tetranucleosome. This aggregation resulted in contacts between free nucleosomes and those within the tetra-nucleosome. Such contacts can be clearly observed in the end configuration of a typical simulation trajectory provided in Figure 4A. They are also evident in the distributions of free nucleosomes around the tetra-nucleosome. As shown in Figure S28, the radial distributions indicate significant populations at distances as small as 5 nm, a value achievable only with stacked nucleosomes.

**Figure 4:**
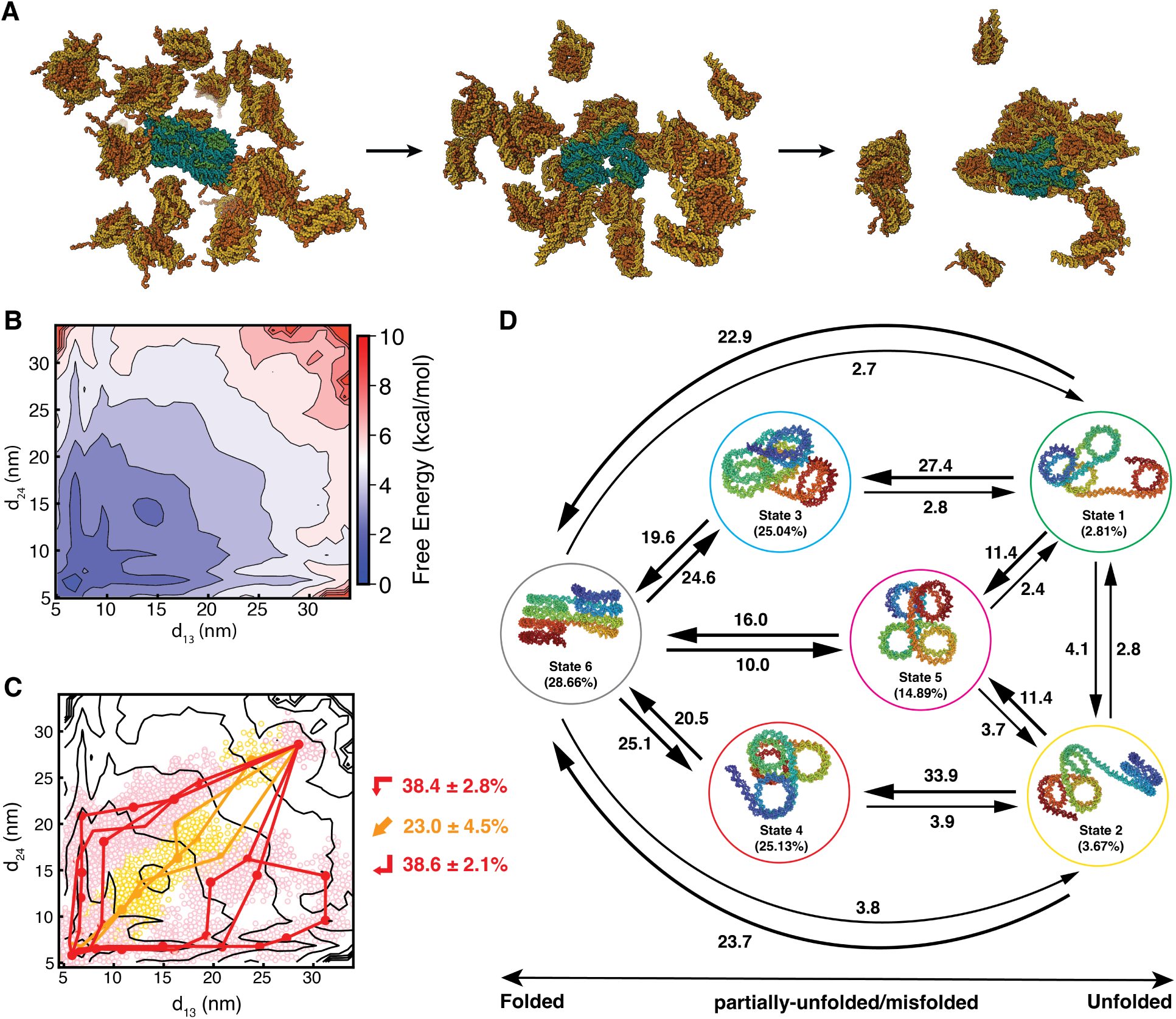
Folding pathways and kinetics for the NRL = 167 tetra-nucleosome embedded in nucleosome condensate. (A) Illustration of the starting, middle, and end configurations of the condensate system along a 70 million step long simulation trajectory. The tetra-nucleosome is shown in cyan and green, and individual nucleosomes are shown in yellow and orange. (B) The free energy profile along the center-of-geometry distance between 1-3 (*d*_13_) and 2-4 (*d*_24_) nucleosomes from the tetra-nucleosome. (C) The three reaction channels for tetra-nucleosome folding. Top three transition pathways with most reactive flux from each one reaction channel are drawn as lines. The filled and open circles represent the centers of, and all the MD configurations belonging to, the microstates along the pathways, respectively. The total reactive flux of each reaction channel is provided on the side. (D) Diagram of the IGME model with six macrostates. The numbers represent rates estimated as inverse MFPT labeled in unit of (10^9^ steps)*^−^*^1^ for the corresponding transitions. Histone proteins are not shown in the representative configurations of each state.

We integrated individual unbiased simulations to construct microstate-MSM for the tetranucleosome in the condensate. By using the equilibrium populations of microstates, we estimated the two-dimensional free energy landscape of the tetra-nucleosome. As shown in Figure 4B, the overall shape of the free energy landscape remains similar to that of the isolated system, but the free energy values of the folding intermediate structures have decreased (see also Figure S29). This reduction is attributed to interactions between the tetra-nucleosome and free nucleosomes, which stabilize partially folded structures. However, the overall change in the free energy landscape is relatively small, on the order of 1-2 kcal/mol. We then identified tetra-nucleosome folding pathways for the condensate system and clustered them into reaction channels. Similar to the isolated system, we found thousands of pathways with comparable fluxes were categorized into sequential or concerted channel. The distribution of fluxes among these reaction channels also remained largely unchanged, indicating a conservation of the folding mechanism. (Figures 4C and S17). The relatively small changes in the free energy landscape and folding reaction channels prompt questions about the role of the nucleosome condensate in tetra-nucleosome folding.

To more directly and accurately examine chromatin folding kinetics, we constructed a six-macrostate IGME model (Figure 4D). Due to the complex interactions introduced by nucleosome condensation, we found that the required lag time to build a Markovian MSM is significantly prolonged (Figure S25). With limited-length MD simulations, the IGME model shows distinct advantages, capturing the slow dynamics with much less error (Figure S24-S25). Based on the optimal IGME model, we observed an obvious increase in the populations of partially unfolded states 3 and 4, which are comparable to the population of the folded state 6. This finding underscores the changes in the free energy observed in Figure 4B and the impact of nucleosome condensate on chromatin stability. Notably, the unfolding rates from state 6 to these intermediate states are faster than the folding rates. Therefore, the nucleosome condensate does not fundamentally alter the folding modes of the tetra-nucleosome but quantitatively modulates the free energy landscape, favoring unfolding dynamics driven by inter-nucleosomal contacts.

### The role of DNA linker length on chromatin folding

Our study thus far has focused on a tetra-nucleosome with an NRL of 167 bp (i.e., 20-bp linkers). However, linker length is known to influence chromatin organization significantly.^40,64–68^ A DNA linker of 10*n* bp facilitates well-aligned stacking between *i* and *i* + 2 nucleosomes to form the zigzag fibril structure.^6,7,69,70^ Extending the linker by 5 bp induces a half-turn twist in the DNA, disrupting the stacking between *i* and *i* + 2 nucleosomes and destabilizing the zigzag conformation.^71^ These 10*n* + 5 linkers are significant and have been observed in certain mammalian cells.^72^ However, structural characterization of chromatin with 10*n* + 5 linker DNA is scarce due to the irregularity and instability of their organization. We further investigate how increasing the linker by 5 and 10 bp impacts chromatin stability and folding dynamics using the computational pipelines mentioned earlier.

The folding of the tetra-nucleosome with an NRL of 177 resembles that of an NRL of 167 but exhibits quantitative differences in reactive fluxes of folding. Its free energy landscape also shows a global minimum at small values of *d*_13_ and *d*_24_ (Figure S26A), indicating stacked two-column structures. However, the stability of this folded state is reduced, as evidenced by its lower equilibrium populations in the six-macrostate IGME model (Figure S26B). The folded conformations bring linker DNA into close proximity, increasing electrostatic repulsion for longer linkers, which decreases stability. Intermediate states along the sequential pathway also feature stacked nucleosomes with closely positioned linker DNA, resulting in similar electrostatic penalties. As a result, unlike the NRL = 167 tetra-nucleosome, the NRL = 177 one favors the concerted folding pathway over the sequential ones (Figure S17D). Notably, our findings align with those of Correll et al. ^65^, who observed an expansion of chromatin with longer 10*n*-bp linkers.

Unlike systems with 10*n*-bp linker DNA, the NRL = 172 tetra-nucleosome system explores distinct chromatin conformations over a different folding landscape. As shown in Figure 5A, the free energy profile has a broader global minimum and a smaller free energy gap between the folded and unfolded states. This broad basin is due to the absence of a single stable conformation. Instead, an ensemble of irregular structures are equally probable. The six-macrostate IGME model (Figure 5B) reveals that two states, 5 and 6, emerge as “folded” states with comparable populations. Unlike the folded states in 10*n* bp linker systems, these states do not feature a dominant stable configuration but rather an ensemble of conformations (Figures 5C and 5D). While the structures in states 5 and 6 are compact and promote protein-DNA interactions, they do not exhibit perfect stacking between *i* and *i* + 2 nucleosomes due to the topological constraints imposed by the additional 5 bp. Instead, they resemble intermediate states observed along the folding pathways for 10*n* bp linker systems. The high transition rates between states 5 and 6 further emphasize the dynamic nature of the tetra-nucleosome with 10*n* + 5 bp linkers.

**Figure 5:**
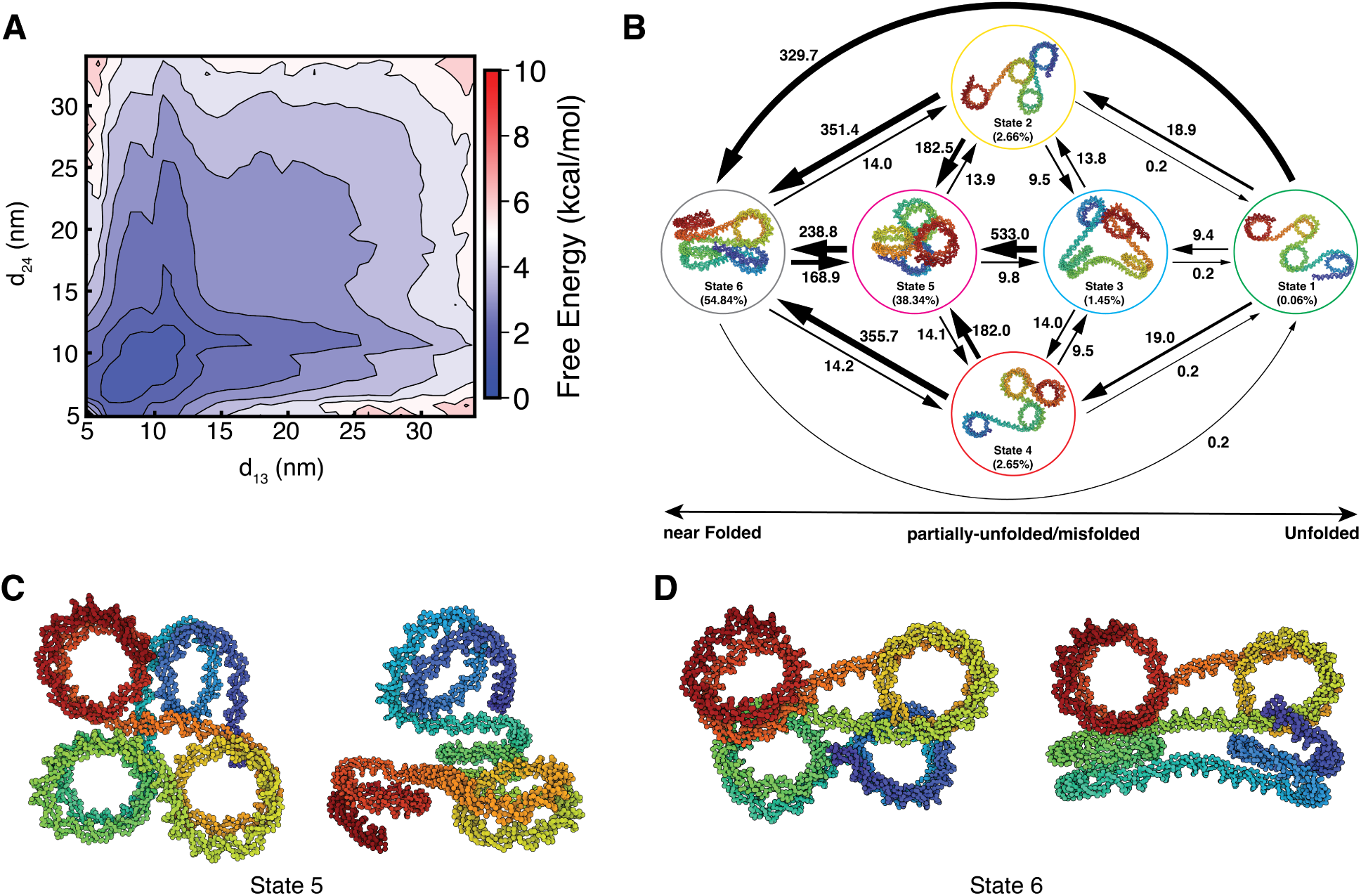
Folding pathways and kinetics for the NRL = 172 tetra-nucleosome. (A) The free energy profile along the center-of-geometry distance between 1-3 (*d*_13_) and 2-4 (*d*_24_) nucleosomes from the tetra-nucleosome. (B) Diagram of the IGME model with six macrostates. The numbers represent rates estimated as inverse MFPT labeled in unit of (10^9^ steps)*^−^*^1^ for the corresponding transitions. Histone proteins are not shown in the representative configurations of each state. (C,D) Addition representative structures the compact stable states 5 and 6.

The unfolded states, 1, 2, and 4, which have more extended structures in the NRL = 172 tetra-nucleosome, also reveal distinct topology. Unlike the extended states seen in 10*n* bp linkers, where consecutive nucleosomes reside on the same side of the linker DNA, the half-turn linker DNA in NRL = 172 tetra-nucleosome introduces additional twist, causing consecutive nucleosomes to reside on opposite sides of the linker.

## Conclusions and Discussion

We combined CG MD simulations, Markov state modeling, and non-Markovian dynamic modeling techniques to comprehensively characterize chromatin stability and folding mechanisms. The implementation of the non-Markovian IGME model allows us to accurately investigate long-timescale folding dynamics, which are challenging to capture through direct simulations or conventional MSMs. Our study of a single tetra-nucleosome reveals multiple folding pathways, with intermediate states resembling *in vivo* chromatin organization.^15,73,74^ For instance, intermediates along the concerted pathway often adopt a coplanar geometry, with all four nucleosomes in the same plane, resembling the *β*-rhombus configuration. ^15^ In contrast, intermediates along the sequential pathway have at least one nucleosome out of the plane, similar to the *α*-tetrahedron structure.^15^ These findings suggest that the absence of regular fibril configurations inside cell nuclei may result from chromatin unfolding driven by factors such as the condensate environment and linker length.

Our examination of the tetra-nucleosome within the nucleosome condensate supports the hypothesis that a crowded environment, typical inside cell nuclei, can drive chromatin unfolding.^18^ The stacking interactions that stabilize the fibril configuration in a single chromatin chain may be replaced by contacts between nucleosomes from different chains. The replacement leads to the formation of irregular conformations, even for DNA linker length that favor fibiril structures.

Additionally, we found that 10*n*+5 bp linkers destabilize the zigzag fibril configurations, favoring structures that resemble *in vivo* configurations observed by cryo-electron tomography.^73,74^ Chromatin with these DNA linkers tends to adopt compact conformations without perfect stacking between *i* and *i* + 2 nucleosomes. These conformations are irregular and form a dynamic ensemble, similar to the behavior of intrinsically disordered proteins.

Our study demonstrates that residue-level CG models enable predictive modeling of chromatin organization at near-atomistic resolution, particularly in conditions challenging for experimental techniques, while the novel non-Markovian IGME models facilitate in-depth qualitative analysis of long-term chromatin conformational dynamics. Although our focus was on the impact of nucleosome condensate on chromatin organization, our approach can be generalized to study other protein condensates, such as HP1*α* for constitutive heterochromatin. Recent advancements in force fields and software highlight the potential for exciting future research directions.^61,75,76^

## Methods

### Coarse-grained molecular dynamics simulations of chromatin

Following previous studies,^17,18^ we utilized residue-level CG models to examine chromatin conformational dynamics. Specifically, we combined the 3SPN.2C model,^48,77,78^ which represents DNA with three sites per nucleotide, and the SMOG model,^79^ which represents histone proteins with one bead per amino acid, to model chromatin. Protein-DNA interactions included electrostatic interactions and non-specific Lennard-Jones potentials. Electrostatic interactions are calculated at 300 K and 150 mM ionic strength using the Debye-Hückel potential. This setup has been shown to replicate the energetics of DNA unwinding in single nucleosomes^60^ and the force-extension curves of chromatin fibers.^18^ To reduce computational cost and prevent nucleosome sliding, we rigidified the histone core and the inner 73 bp layer of core DNA during all simulations. Ding et al. ^17^ demonstrated that such rigidification has negligible effects on the chromatin folding landscape.

We studied three isolated tetra-nucleosome systems with NRL of 167, 172, and 177 bp, respectively. For each system, we performed 4,643 independent, unbiased MD simulations starting from distinct initial configurations. Initial configurations for these simulations were prepared using 10,000 tetra-nucleosome structures from a previous study that applied enhanced sampling methods to comprehensively cover the configuration space.^17^ We computed the inter-nucleosome distances for the 10,000 structures and selected those with *d*_13_ ≥ *d*_24_ (4,643 in total) to serve as reference values for further MD simulations. From these reference inter-nucleosome distances, we generated the initial configurations using 0.2 million-step-long restrained MD simulations to bias the *d*_13_ and *d*_24_ toward reference values. From these initial configurations, we conducted unbiased NVT simulations for production. For the NRL = 167 system, each trajectory lasted at least 0.8 million steps, while for the NRL = 172 and NRL = 177 systems, the trajectory length was consistently set to 1 million steps. All simulations were performed at 300 K using LAMMPS on CPUs,^80^ and the pairwise distances between each pair of nucleosome geometric centers (i.e., (*d*_12_*, d*_13_*, d*_14_*, d*_23_*, d*_24_*, d*_34_)) were recorded every 500 steps.

Additionally, we conducted simulations for a system where the NRL = 167 tetra-nucleosome is embedded in a nucleosome solution. Due to the higher computational cost of this system, we limited the number of independent simulations to 530. Initial configurations of the tetra-nucleosome for these simulations were prepared by selecting structures from the centers of the 530 microstates constructed for the isolated NRL = 167 tetra-nucleosome. Then we placed each of the tetra-nucleosomes in the center of a cubic box with a 55 nm edge, and randomly placed 26 additional single nucleosomes to achieve a total nucleosome concentration of 0.3 mM. We relaxed each of these initial configurations with a 0.2 million-step NVT simulation, during which the tetra-nucleosome was fixed as a rigid body while single nucleosomes were free to move. From the relaxed configurations, we conducted production NVT simulations of at least 7 million steps, allowing all nucleosomes to evolve freely. Simulations were performed at 300 K with OpenMM and OpenABC packages on GPUs.^76,81^ The pairwise distances between nucleosomes within the tetra-nucleosome, as well as the positions of all nucleosomes, were recorded every 500 steps for analysis.

Additional details on the system setup and MD simulations are provided in the *Supporting Information*.

### Markov state modeling and transition path analysis for chromatin folding

We constructed independent microstate-MSMs and performed transition path analysis for all four systems following a consistent protocol, briefly outlined below. Additional details are provided in the *Supporting Information*.

a. Select the converged segments of the trajectories and duplicate the converged trajectories according to the reflection symmetry of the nucleosome indices. All the following analyses are applied to the converged and duplicated trajectories.
b. With six inter-nucleosomal distances *d_ij_* as input features, apply the time-lagged independent component analysis (tICA) method with kinetic mapping algorithm to further reduce dimensionality and uncover independent collective variables (CVs) that better represent slow timescale dynamics.^55–57,82^
c. Group MD conformations into microstates by the K-Means algorithm according to their kinetic similarities based on the geometric distances in the tICA CV space. The hyperparameters of the tICA and clustering (i.e., tICA relaxation time, number of tICs, and number of microstates) are optimized using the cross-validation with the generalized matrix Rayleigh quotient (GMRQ) score.^83^
d. Construct and validate the microstate-MSMs by the Chapman-Kolmogorov (CK) test and implied time scale (ITS) analysis.^25,26^
e. Employ the Transition Path Theory (TPT)^20,21,58^ to elucidate the folding kinetic pathways from unfolded states to folded states and their corresponding fluxes.
f. Lump multiple parallel kinetic pathways into a small set of metastable and representative reaction channels using Latent-space Path Clustering (LPC) algorithm to facilitate the understanding of folding mechanisms. ^27,32,84^

All analyses were conducted using in-house Python codes and codes based on MSM-Builder version 3.8.1.^85^

### IGME modeling of chromatin folding

To facilitate the interpretation of chromatin folding, for each simulation system, we modeled its folding dynamics with a six-macrostates IGME model ^36^ (Figure 2D). These six macrostates were constructed by lumping hundreds of microstates using the Robust Perron Cluster Analysis algorithm.^86,87^ The IGME model encodes non-Markovian dynamics through time integrations of memory kernels based on the GME. ^36^ Specifically, IGME model describes the evolution of transition probabilities over time using **T**(*t* ≥ *τ_k_*) = **A**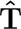*^t^*, where **T**(*t*) is the transition probability matrix (TPM) at lag time *t*, and *τ_k_* is the relaxation time of the memory kernels. The matrices **A** and 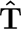 are constant matrices estimated from the MD simulation data.

We determined the relaxation time of the memory kernels, *τ_k_*, by examining the convergence of the mean integral memory kernels. This can be quantified using two approaches: the first involves our previously developed quasi-MSM (qMSM)^38^ to compute discretized kernels across various lag times, while the second directly utilizes the IGME to approximate the time-integrated memory kernels.^36^ The **A** and 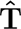 matrices were estimated using a set of TPMs at short lag times through least squares fitting with a Lagrangian approach. The optimal fitting range was selected by a systematic scanning to minimize the deviation between the predictions from IGEM model and raw MD data. The optimal IGME models were further validated and compared to MSMs using the Chapman-Kolmogorov test. All reported thermodynamic and kinetic properties of the metastable macrostates were derived from the optimal IGME models, which exhibit the smallest deviations from the raw MD data. More details and results about IGME models are presented in the *Supporting Information*.

## Acknowledgement

This work was supported by the National Institutes of Health (Grant R35GM133580) and the National Science Foundation (Grant MCB-2042362). X.H. acknowledge the support from the Hirschfelder Professorship Fund from University of Wisconsin-Madison and the Research Forward Fund from the University of Wisconsin-Madison Office of the Vice Chancellor for Research with funding from the Wisconsin Alumni Research Foundation. The computational resource support from Center for High Throughput Computing at the University of Wisconsin-Madison is also acknowledged.^88^

## Data and materials availability

Data presented in this study is available upon reasonable request to the corresponding author.

